# Evidence of a Causal Association Between Cancer and Alzheimer’s Disease: a Mendelian Randomization Analysis

**DOI:** 10.1101/653352

**Authors:** Sahba Seddighi, Alexander L Houck, James B Rowe, Paul DP Pharoah

**Affiliations:** University of Cambridge Institute of Public Health, University of Cambridge, Cambridge, UK; University of Tennessee Health Science Center, Memphis, Tennessee, US; Department of Clinical Neurosciences, University of Cambridge, Cambridge, UK; Department of Oncology, Cambridge University Hospitals NHS Foundation Trust, Cambridge, UK

## Abstract

**Objectives:** To determine whether cancer confers protection against Alzheimer’s disease and to evaluate the relationship in the context of smoking-related cancers versus non-smoking related cancers

**Design:** Mendelian randomization analysis using cancer-associated genetic variants as instrumental variables

**Setting:** International Genomics of Alzheimer’s Project

**Participants:** 17,008 Alzheimer’s disease cases and 37,154 controls

**Main outcome measures:** Odds ratio of Alzheimer’s disease per 1-unit higher log odds of genetically predicted cancer

**Results:** We found that genetically predicted lung cancer (OR 0.91, 95% CI 0.84-0.99, p=0.019), leukemia (OR 0.98, 95% CI 0.96-0.995, p=0.012), and breast cancer (OR 0.94, 95% CI 0.89-0.99, p=0.028) were associated with 9.0%, 2.4%, and 5.9% lower odds of Alzheimer’s disease, respectively, per 1-unit higher log odds of cancer. When genetic predictors of all cancers were pooled, cancer was associated with 2.5% lower odds of Alzheimer’s disease (OR 0.98, 95% CI 0.96-0.988, p=0.00027) per 1-unit higher log odds of cancer. Finally, genetically predicted smoking-related cancers showed a more robust inverse association with Alzheimer’s disease than non-smoking related cancers (5.2% lower odds, OR 0.95, 95% CI 0.92-0.98, p=0.0026, vs. 1.9% lower odds, OR 0.98, 95% CI 0.97-0.995, p=0.0091).

**Conclusions:** Genetically predicted lung cancer, leukemia, breast cancer, and all cancers in aggregate are associated with lower odds of incident Alzheimer’s disease. Furthermore, the risk of Alzheimer’s disease was lower in smoking-related versus non-smoking related cancers. These results add to the substantial epidemiological evidence of an inverse association between history of cancer and lower odds of Alzheimer’s disease, by suggesting a causal basis for this relationship.

## 1. Introduction

Individuals with a history of cancer are less likely to develop Alzheimer’s disease (AD), and vice versa.^1–3^ This epidemiological observation remains unexplained, although biologically plausible mechanisms have been proposed: while cancer is a disorder of excessive cell proliferation, neurodegeneration is one of premature cell death. The dysregulation of mutual genes, proteins, and pathways in both conditions has been demonstrated.^4^ For instance, the expression of the tumor suppressor TP53, which is notably downregulated in the majority of cancers, is upregulated in AD. The opposite is true for PIN1, an enzyme promoter of cell proliferation.^4^

The magnitude of the association between AD and cancer varies greatly across studies. In one population-based prospective study, those with prevalent cancer had a 69% lower risk of developing AD after controlling for numerous factors.^5^ Another found that those with a history of cancer had just a 13% lower risk of subsequent AD.^6^ Others report a trend without reaching statistical significance,^7 8^ or no association between a history of cancer and the risk of AD.^9^ In light of these findings, it is important to consider the potential for bias and confounds behind these associations. The reduced risk of AD in those diagnosed with cancer may be due to higher mortality rates among cancer patients and survivors, especially for cancers with poor survival rates, such as pancreatic cancer.^10^ Many studies are limited by small sample sizes^7 11 12^, short follow-up periods^7 9 12 13^, or gender bias.^9^ Other issues include appropriate diagnoses and changing guidelines, the potential negative effects of chemotherapy on cognition, and behavioral and sociodemographic variables related to both AD and cancer.

A fundamental question remains: is cancer causally related to AD, or is the observed epidemiological relationship between the two disorders an artifact of study design, confounds and biases? This study assesses potential genetic mechanisms by which cancer confers protection against subsequent AD, by using a Mendelian randomization approach. Prior literature was used to identify genetic variants that influence susceptibility to cancer. Data from the International Genomics of Alzheimer’s Project (IGAP) were then utilized to examine the theoretically unconfounded association between these cancer-predicting genes and the risk of AD. This approach addresses two major limitations of previous observational studies: the potential for reverse-causation and the confounding effects of environmental risks.^14^ We predicted that genes that have been associated with increased risk of cancer are associated with reduced risk of AD, in independent cohorts.

## 2. Methods

### Overview

The summary-data-based Mendelian randomization method consists of three key steps in data preparation and cleaning: (1) the identification of single nucleotide polymorphisms (SNPs) associated with the exposure (in this case, cancer), (2) the determination of SNP-effects on the outcome (in this case, AD), and (3) Mendelian randomization analysis to examine the causal effect of the exposure on the outcome (Figure 1).

**Figure 1.**
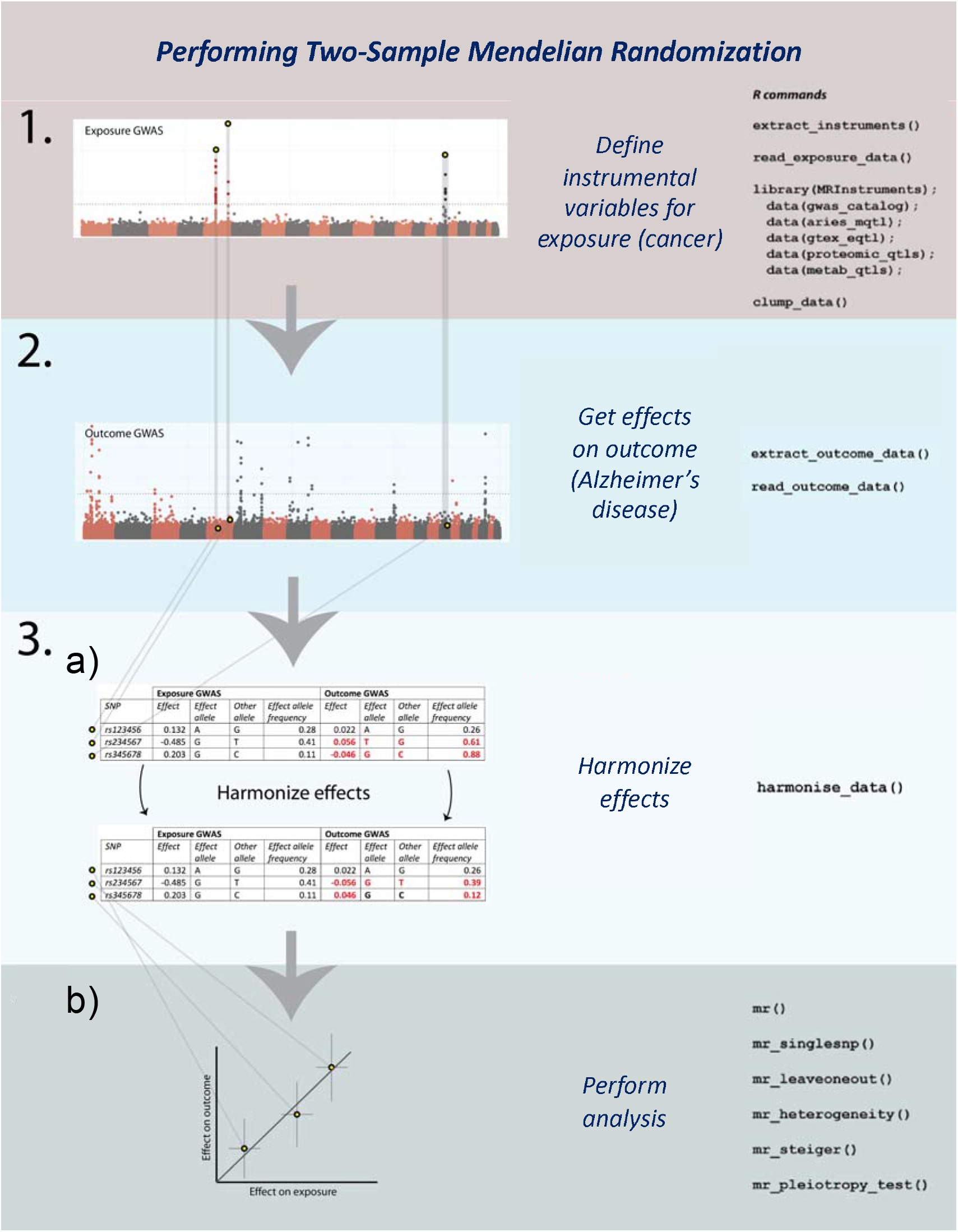
Schematic overview of steps involved in two-sample Mendelian randomization. 1) After the identification of relevant cancer types, genetic variants associated with each cancer type (i.e. instrumental variables) were identified from published GWAS. 2) Summary statistics for the instrumental variables were extracted from the International Genomics of Alzheimer’s Project, the outcome GWAS. 3) Mendelian randomization analysis was performed by a) harmonizing the exposure and outcome summary statistics files and b) calculating an estimate for the causal effect of cancer-associated SNPs on risk of AD.

### 2.1 Defining instrumental variables for cancer

#### 2.1.1 Identifying relevant cancer types: systematic literature review

To identify cancer types with epidemiological evidence of association with AD, PubMed (NCBI) and Web of Science (EBSCO) were systematically searched using a combination of medical subject headings [MeSH] and keyword strings (Box 1).

##### Box 1: Literature Search Strategy

**Pubmed (NCBI): searched on 04/06/18**

**Step 1:** neoplasms [MeSH Terms] OR cancer [All Fields]

**Step 2:** Alzheimer disease [MeSH terms] OR Alzheimer* [All Fields]

**Step 3:** Epidemiological studies [MeSH terms] OR cohort studies [MeSH terms] OR prospective studies [MeSH terms] OR longitudinal [All Fields] OR follow-up [All Fields] OR observational study [All Fields]

**Search:** 1^ AND 2^ AND 3^

**Filters:** Full text; humans

**Web of Science (EBSCO): searched on 04/06/18**

**Step 1:** Cancer (Topic)

**Step 2:** Alzheimer disease (Topic) Step3: Epidemiological study (Topic)

**Search:** 1^ AND 2^ AND 3^

**Filters:** English; open access

Identified papers were then examined against a predetermined set of inclusion criteria to determine eligibility (Box 2).

##### Box 2: Inclusion Criteria

- Full text
- Published in English
- Observational cohort studies recruited at the population level (i.e. not hospital-based)
- Longitudinal design
- Cancer diagnosis precedes AD diagnosis
- Incidence of AD can be compared between those with and without cancer

#### 2.1.2 Identifying cancer-associated genetic variants: review of GWAS literature

After identifying cancers with prior evidence of association with AD, the Genome-Wide Association Studies (GWAS) Catalog was used to identify genetic variants associated with each cancer type.^15^ The GWAS Catalog (https://www.ebi.ac.uk/gwas/docs/about) was launched in 2008 by the National Human Genome Research Institute to systematically catalogue and summarize SNP-trait associations from all published GWAS.^15^ The repository is updated on a weekly basis by an experienced team of molecular biologists with support from The European Bioinformatics Institute (EMBL-EBI) and input from an independent Scientific Advisory Board. This tool was used to identify published GWAS reporting cancer susceptibility SNPs. Each paper was then manually examined for content and data retrieval.

Only studies conducted or replicated in individuals of European decent were considered in order to enable meaningful comparison with the population used in IGAP for Mendelian randomization outcome data. Furthermore, studies conducted on susceptible subpopulations (e.g. those with a family history of cancer, or ever-smokers) were not considered. SNPs associated with cancer at the standard threshold for genome-wide significance (p<5×10^−8^) were selected.

A central assumption of Mendelian randomization is that instrumental variables are free of linkage disequilibrium with one another (i.e. each genetic variant must be inherited independently of all others under consideration). Analyses were, therefore, limited to genetic variants that were not in linkage disequilibrium (defined as r^2^<0.2) with other genetic variants for the same type of cancer in order to avoid a violation of this assumption. Pairwise linkage analysis of SNPs was undertaken using the National Cancer Institute LDMatrix tool (https://analysistools.nci.nih.gov/LDlink/?tab=ldmatrix) based on the European subpopulation—Utah Residents from North and West Europe (CEU), Toscani in Italia (TSI), British in England and Scotland (GBI), and Iberian population in Spain (IBS)—reference panel of the 1000 Genomes Project. For genetic variants in linkage disequilibrium, the variant with the lowest p-value for association with cancer was selected. Similarly, for each cancer susceptibility locus, the lead SNP, representing the variant with the lowest p-value of association, was selected.

#### 2.1.3 Identifying proxy SNPs

When cancer-associated genetic variants were unavailable in the outcome dataset, proxies in pairwise linkage disequilibrium (defined by r^2^ >0.9) were used, where available. Identification of proxy SNPs was undertaken using the National Cancer Institute LDlink platform’s LDproxy tool (https://analysistools.nci.nih.gov/LDlink/?tab=ldproxy) based on the European subpopulation— CEU, TSI, GBI, and IBS— reference panel of the 1000 Genomes Project. Proxy SNP effect alleles were assigned according to correlation information between alleles, provided on the LDpair tool provided through the National Cancer Institute LDlink platform.

### 2.2 Obtaining effects of cancer-associated SNPs on AD

Summary statistics describing the association between each cancer-related SNP and risk of AD were obtained from IGAP.^16^ IGAP includes genotyped and imputed data on SNPs from 17,008 AD cases and 37,154 controls of European ancestry from four genome-wide association studies: (1) The Cohorts for Heart and Aging Research in Genomic Epidemiology consortium (CHARGE), (2) The Alzheimer’s Disease Genetics Consortium (ADGC), (3) The Genetic and Environmental Risk in Alzheimer’s disease consortium (GERAD), and (4) The European Alzheimer’s disease Initiative (EADI). A summary of each dataset is provided in Supplementary Table 1. Diagnostic criteria for AD used in each study are provided in Supplementary Table 2.

#### 2.2.1 Imputation and SNP selection

SNPs with call rates <95% were excluded from consideration in IGAP. The genotypes of all individuals were imputed with haplotypes from samples of European ancestry in the 1000 Genome Project using IMPUTE2^17^ or MaCH/Minimac.^18^ SNPs with a minor allele frequency of <1% or with R^2^ (MaCH) or info score quality (IMPUTE2) estimates less than 0.3 were excluded from analyses. A total of 8,133,148 SNPs was retained for analysis.

In each dataset, the association between genotype dosage and AD was analyzed by logistic regression. The model was adjusted for age, sex, and principal components to account for possible population stratification. For the CHARGE cohorts containing incident AD cases, Cox proportional hazards regression models were used instead. The consortia used different, but comparable software for these analyses (PLINK (ACT, ADC, ADNI, AGES, GSK, MAYO, OHSU, ROSMAP, TGEN2, UMVUMSS, and UPITT), SNPTEST (GERAD), ProbABEL (AGES, CHS, EADI, FHS, and RS) or R (LOAD, MIRAGE).^19–21^ SNPs with logistic regression |β| > 5 or p-value equal to 0 or 1 were excluded. The maximum number of SNPs in any data set was 8,131,643. SNPs that were genotyped/imputed in at least 40% of AD cases and 40% of controls were included in the meta-analysis. This led to a final number of 7,055,881 SNPs for analysis.

#### 2.2.2 Statistical analysis

For the meta-analysis, fixed-effects inverse variance-weighted meta-analysis was applied, with the standard errors of the β-coefficient scaled by the square roots of study-specific genomic inflation factors (estimated before combining summary statistics across all datasets). METAL and GWAMA software packages were used for these analyses.^22 23^

### 2.3 Statistical analysis of two-sample Mendelian randomization

#### 2.3.1 Calculating inverse-variance weighted ratio estimates and evaluating the role of chance

Two-sample Mendelian randomization is a statistical method that can be applied to summary statistics from GWAS to estimate the causal effect of an exposure (in this case, cancer) on an outcome (in this case, AD). “Two-sample” refers to the fact that summary association results for the exposure and the outcome are estimated in non-overlapping sets of individuals.^24^

An instrumental variable ratio estimate was calculated for each cancer-associated SNP as follows: First, the exposure and outcome files were harmonized on SNPid and effect allele. Then, the effect size estimate (β) for the association of the SNP with risk of AD was divided by the effect size estimate for the association of the same SNP with risk of cancer, producing a ratio of association estimates for each 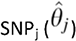 as follows^25^:

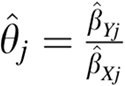

The ratio estimates from each SNP were then averaged using the inverse-variance weighted formula, adopted from the meta-analysis literature, to produce an overall causal estimate, the inverse-variance weighted (IVW) estimate. The variance term was calculated as^25^:

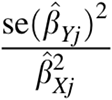

And the pooled fixed-effect inverse-variance weighted estimate 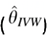 was calculated as^25^:

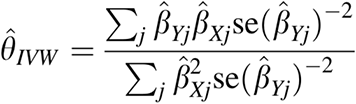

Ratio estimates for each genetic variant were summarized as forest plots. All analyses were conducted in R version 3.5.0 (R foundation) using the two sample Mendelian Randomization package. The full R script used for two-sample MR is provided in Appendix 1.

To assess the role of chance in our findings, we additionally calculated the Bayesian false-discovery rate (BFDR) for each result, irrespective of statistical significance. Under frequentist inference, p-values represent the probability of the data given the hypothesis. A Bayesian approach can be used, instead, to assess the probability of a hypothesis given the data. BFDR can be calculated to identify noteworthy associations and prevent overinterpretation of statistically significant findings that are unlikely to be true. This method involves specification of the prior probability that the null hypothesis is false and combines this information with the p-value and study power to derive a more informative posterior probability that the null hypothesis is true. A range of priors was selected on the basis of available observational evidence. All analyses were conducted in R version 3.5.0 (R foundation) using the gap package. The full R script used for BFDR calculations is provided in Appendix 1.

#### 2.3.2 Sensitivity analyses

##### 2.3.2.1 MR-Egger intercept test

Conventional Mendelian randomization analysis relies on the assumption that genetic variants used as instrumental variables do not have pleiotropic effects—meaning that the chosen variants do not influence the outcome (AD) through any pathways other than the exposure (cancer). This, therefore, implies that the only causal pathway from a cancer-associated genetic variant to AD is via cancer associated biological pathways. While this may be a reasonable assumption when the risk factor under study is a protein biomarker and SNPs are located in or near the coding region for that protein, it is more questionable for polygenic exposures, such as cancer.^25^ We, therefore, used the MR-Egger intercept test to detect violations of this assumption, assessing whether cancer-associated genetic variants have pleiotropic effects on AD that differ on average from zero— known as directional pleiotropy.^25^ If the intercept is not significantly different from 0, there is no evidence to reject the null hypothesis of no directional pleiotropy. MR-Egger intercepts were generated via the two-sample Mendelian randomization package, using R script provided in Appendix 1.

##### 2.3.2.2 Leave-one-out analysis

To assess the robustness of the Mendelian randomization effect estimates and identify any potential outliers, each SNP was sequentially removed from analysis, conducting Y analyses with Y-1 datapoints^25^. If the precision and direction of association between cancer-predicting SNPs and AD remain largely unaltered, this implies that the data are not driven by any outliers. Leave-one-out analyses and corresponding plots were generated using the two-sample Mendelian randomization package via R script provided in Appendix 1.

##### 2.3.2.3 Funnel plots of inverse standard error

Finally, any heterogeneity of effect estimates was visualized using funnel plots depicting the causal effect estimates for each SNP on the x-axis and the inverse standard error (a measure of instrumental strength) for the association on the y-axis. Asymmetry about the vertical line is indicative of heterogeneity; furthermore, a correlation between effect size and instrument strength (which would result in asymmetry) is an indicator of possible directional pleiotropy.^26^ The funnel plots were generated using the two-sample Mendelian randomization package via R script provided in Appendix 1.

### Patient involvement

No patients were involved in the design, recruitment, or conduct of this study. No patients were asked to advise on the interpretation of results or the writing of manuscript. Following publication, these results will be available to the general public.

## 3. Results

### 3.1 Defining instrumental variables for cancer

#### 3.1.1 Identifying cancer types of interest

Following a systematic review of titles, abstracts, and content for eligibility, eleven papers were retrieved from PubMed and Web of Science (Figure 2).^5–9 11 13 27–30^ Searching their respective bibliographies did not lead to any additional studies for inclusion. The results of the systematic literature review are summarized in Table 1 and Supplementary Tables 3 and 4.

**Table 1.**
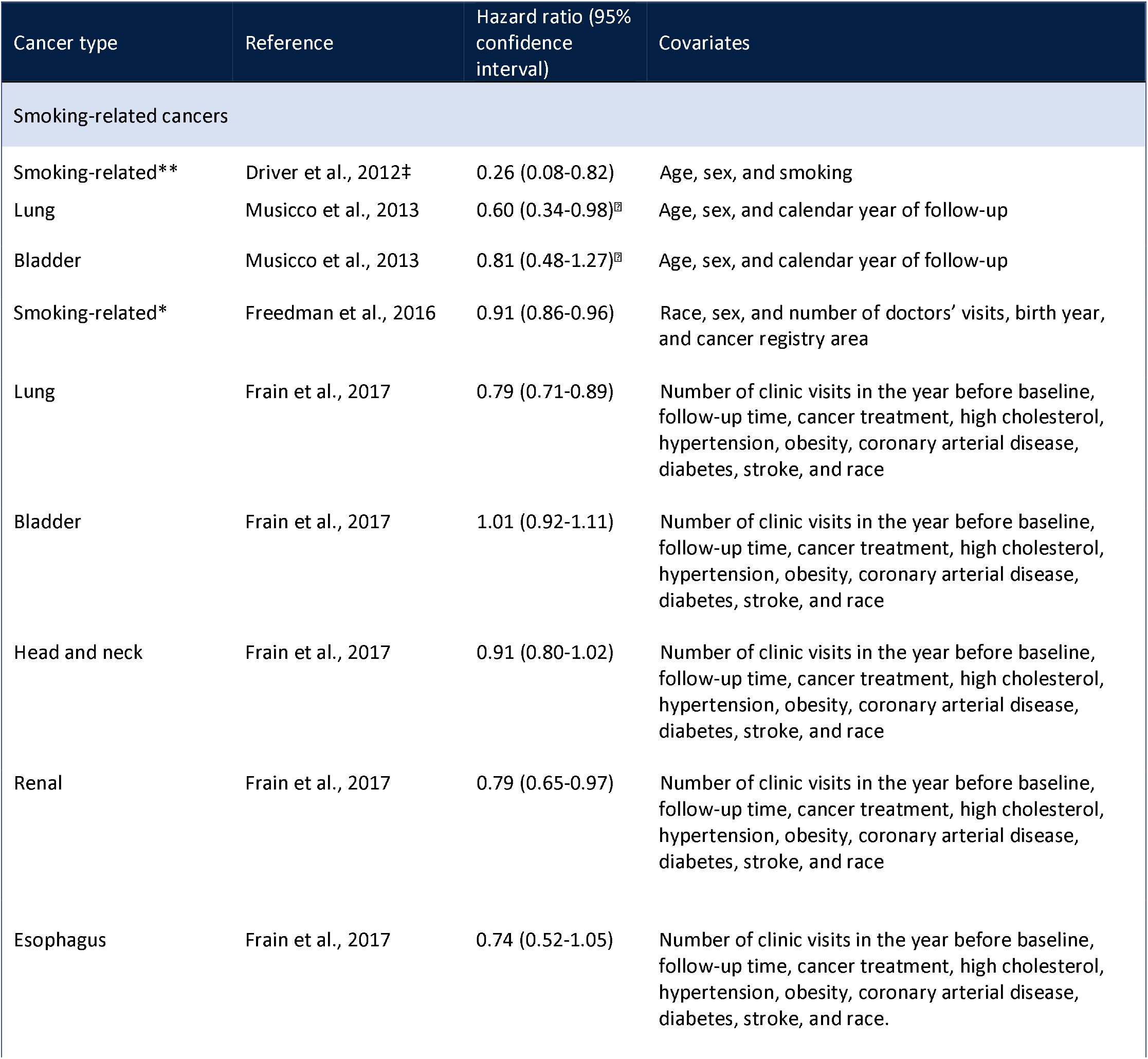
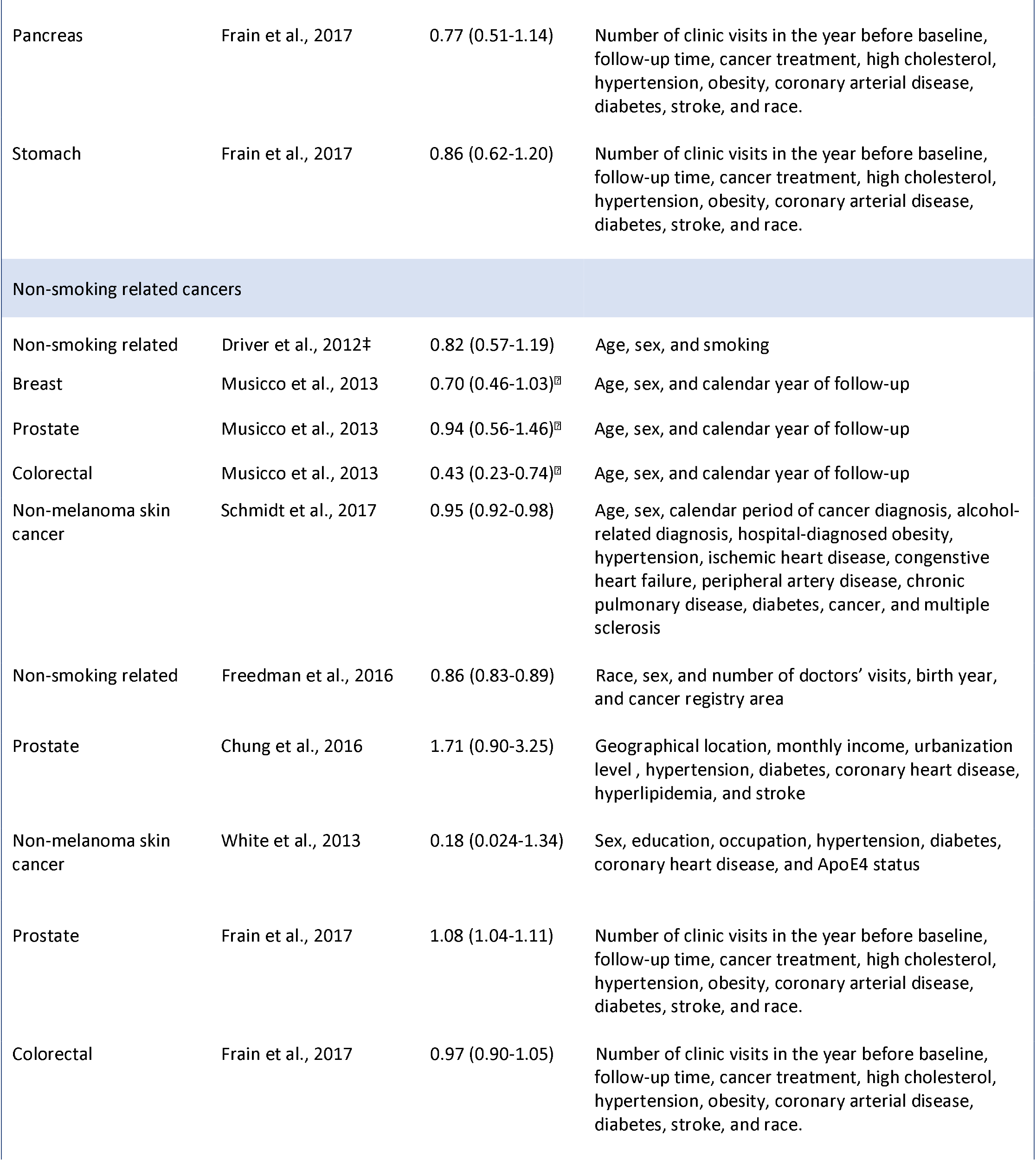
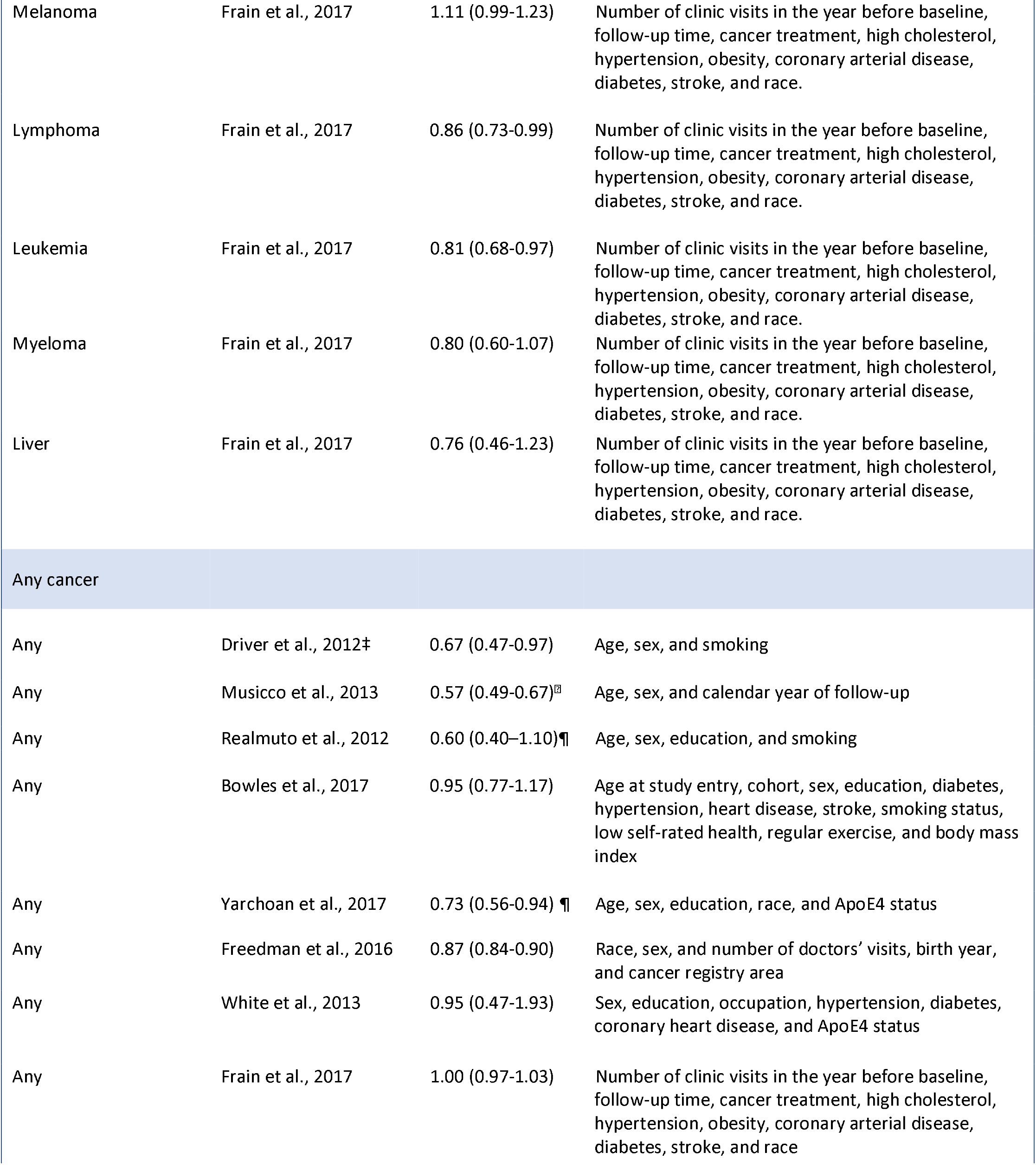
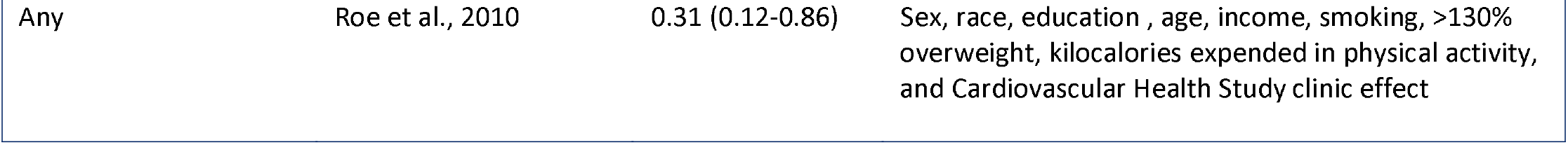
Observational evidence for cancer and risk of subsequent Alzheimer’s disease, by cancer type. Separate effect estimates are provided for smoking-related cancers, non-smoking related cancers, and all cancers together. Unless otherwise noted, the maximally adjusted effect estimate is displayed. * Smoking-related cancers include oral cavity and pharynx, lip, pancreas, lung and bronchus, larynx, cervix, kidney and renal pelvis, bladder, esophagus, and stomach. ** Smoking-related cancers include oral, pharynx, larynx, esophagus, stomach, pancreas, lung, cervix, bladder, and kidney. ‡Maximally adjusted model was not chosen here, due to insufficient statistical power. ⍰ Relative risk ¶ Odds ratio.

**Figure 2.**
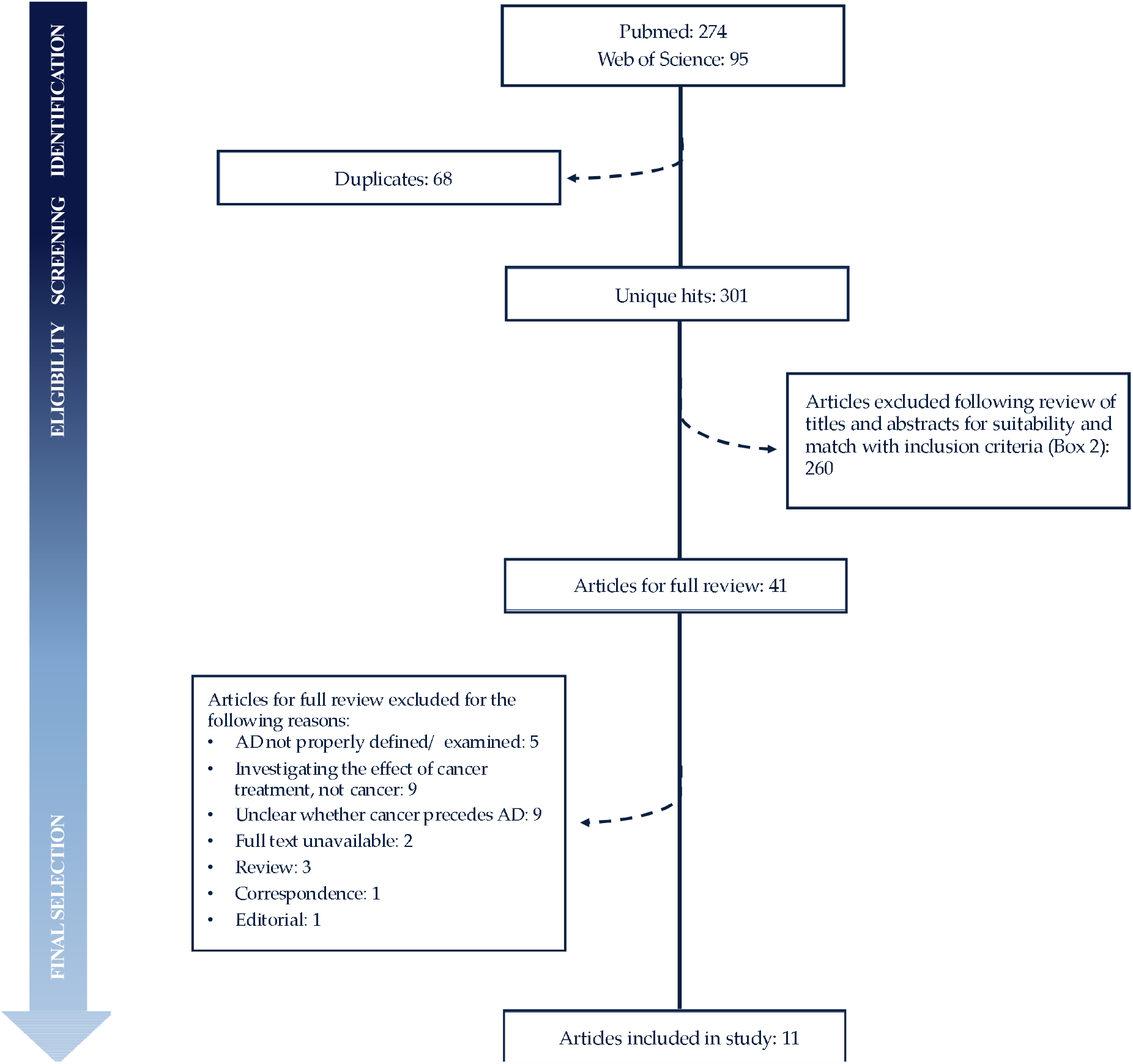
Schematic overview of systematic literature search process and results.

With the exception of two papers, all studies report an inverse association between cancer and risk of subsequent AD, albeit not always statistically significant at standard thresholds. Nominally significant estimates for a protective effect of cancer on risk of subsequent AD range from 13% reduced risk (HR 0.87, 95% CI 0.84 to 0.90)^6^ to 69% reduced risk (HR 0.31, 95% CI 0.12 to 0.86).^5^ The smallest nominally significant estimate for a protective effect of cancer at any individual site on risk of subsequent AD was seen in individuals with non-melanoma skin cancer (HR 0.95, 95% CI 0.92 to 0.98),^27^ while the largest estimate was for colorectal cancer (RR 0.43, 95% CI 0.23 to 0.74).^13^ Only data supporting a protective role of cancer in AD reached statistical significance. However, this inverse relationship is not uniformly observed across individual cancer types. For instance, while one study reports a 57% reduction in risk of AD in those with a history of colorectal cancer (RR 0.43, 95% CI 0.23 to 0.74),^13^ another found no nominally significant difference in the risk of AD in those with a history of colorectal cancer (RR 0.97, 95% CI 0.90 to 1.05). The same is true for non-melanoma skin cancer (HR 0.95, 95% CI 0.92 to 0.98 vs HR 0.18, 95% CI 0.024 to 1.34) (Table 1).^7 27^

Furthermore, Driver and colleagues found a more robust inverse association between smoking-related cancers and AD, as compared to non-smoking related cancers and AD (Table 1). In light of these findings, together with the well-established negative association between smoking and Parkinson’s disease,^31^ another neurodegenerative disorder, subsequent analyses were stratified by smoking versus non-smoking related cancers to further examine this association in the context of AD. Nominally significant effect estimates from observational studies for smoking-related cancers range from 9% reduced risk of AD (HR 0.91, 95% CI 0.86 to 0.96)^6^ to 40% reduced risk of AD (HR 0.6, 95% CI 0.34 to 0.98).^13^ For non-smoking related cancers, nominally significant effect estimates range from 5% reduced risk of AD (HR 0.95, 95% CI 0.92 to 0.98)^27^ to 57% reduced risk of AD (RR 0.43, 95% CI 0.23 to 0.74) (Table 1)^13^.

#### 3.1.2 Identifying cancer-associated genetic variants

GWAS data were available for 6 smoking-related cancers—renal cancer carcinoma, cervical cancer, pancreatic cancer, upper aerodigestive tract cancer, urinary bladder cancer, and lung cancer— in the GWAS Catalog repository. Six additional non-smoking related cancers— prostate cancer, leukemia, breast cancer, melanoma, lymphoma, and ovarian cancer— were selected on the basis of SNP information availability within the repository.

Per the inclusion criteria, 60 GWAS were deemed eligible for consideration (Supplementary Table 5). A total of 314 genetic variants were identified as instrumental variables for cancer (Renal carcinoma: 25; pancreatic cancer: 13; upper aerodigestive tract cancer: 14; urinary bladder cancer: 11; lung cancer: 18; cervical cancer: 1; prostate cancer: 36; leukemia: 38; breast cancer: 109; melanoma: 24; lymphoma: 20; ovarian cancer: 19) (Supplementary Tables 5 and 6). The lead SNP (based on the lowest reported p-value from GWAS) at each locus was selected. The lead SNP was also selected for any SNPs in linkage disequilibrium.

#### 3.1.3 Identifying proxy SNPs

Of the 314 identified genetic variants, 26 were not available in IGAP. Seven SNPs had suitable proxies (defined as r^2^ >0.9) within the European subpopulation reference panel of the 1000 Genomes Project, and nineteen SNPs did not have suitable proxies (Supplementary Table 7).

### 3.2 Obtaining effects of cancer-associated SNPs on Alzheimer’s disease

#### 3.2.1 International Genomics of Alzheimer’s Project

The International Genomics of Alzheimer’s Project datafile consists of the following information for 7,055,881 SNPs and their associations with AD (based on meta-analysis, as described previously): Chromosome (Chromosome of the SNP (Build 37, Assembly Hg19)), Position (position of the SNP (Build 37, Assembly Hg19), MarkerName (SNP rsID or chromosome:position if rsID not available), Effect allele (reference allele (coded allele), Non Effect_allele (non-reference allele (non-coded allele)), Beta-coefficient (overall estimated effect size for the effect allele), Standard error (overall standard error for effect size estimate), P-value (meta-analysis p-value using regression coefficients (beta and standard error))

The IGAP summary statistics corresponding to each cancer-related SNP are detailed in Supplementary Table 6.

### 3.3 Two-sample Mendelian randomization

#### 3.3.1 Inverse-variance weighted effect estimates and false discovery rate

Genetically predicted smoking-related cancers were associated with 5.2% lower odds of AD (OR 0.95, 95% CI 0.92 to 0.98, p=0.0026; BFDR_50%_=7.11%, BFDR_10%_=40.78%, BFDR_1%_= 88.34%, BFDR_0.1%_= 98.71%) per 1-unit higher log odds of cancer (Table 2 and Figure 3). Within smoking-related cancers, only genetically predicted lung cancer, with a 9.0% reduction, was associated with AD (OR 0.91, 95% CI 0.84 to 0.99, p=0.019; BFDR_50%_=20.57%, BFDR_10%_=69.98%, BFDR_1%_= 96.25%, BFDR_0.1%_=99.62%) per 1-unit higher log odds of cancer (Table 2 and Figure 3). As only one susceptibility locus was identified for cervical cancer, separate Mendelian randomization analyses were not conducted for cervical cancer; however, its corresponding SNP was used in the pooled analysis of all smoking-related cancers.

**Table 2.** Results of conventional Mendelian randomization analysis for cancer and Alzheimer’s disease. Genetically predicted lung cancer, leukemia, breast cancer, smoking-related cancers, non-smoking related cancers, and all cancers taken together were associated with significantly lower odds of Alzheimer’s disease. The Bayes False Discovery Rate (BFDR) is also provided for each result as an approximation of the false positive rate. BFDR_50%_= Bayes False Discovery Rate at prior of 50%, BFDR_10%_= Bayes False Discovery Rate at prior of 10%, BFDR_1%_= Bayes False Discovery Rate at prior of 1%, BFDR_0.1%_= Bayes False Discovery Rate at prior of 0.1.

| Cancer type | Odds ratio | Lower confidence limit | Upper confidence limit | P-value | BFDR<sub>50%</sub> | BFDR<sub>10%</sub> | BFDR<sub>1%</sub> | BFDR<sub>0.1%</sub> |
| --- | --- | --- | --- | --- | --- | --- | --- | --- |
| Smoking-related cancers |  |  |  |  |  |  |  |  |
| Renal cell carcinoma | 0.97 | 0.89 | 1.05 | 0.434 | 68.22 | 95.08 | 99.53 | 99.95 |
| Pancreatic cancer | 0.97 | 0.91 | 1.04 | 0.429 | 71.63 | 95.79 | 99.60 | 99.96 |
| Upper aerodigestive tract cancer | 0.96 | 0.90 | 1.03 | 0.235 | 64.60 | 94.26 | 99.45 | 99.95 |
| Urinary bladder cancer | 0.94 | 0.83 | 1.06 | 0.325 | 58.76 | 92.77 | 99.30 | 99.93 |
| Lung cancer | 0.91 | 0.84 | 0.99 | 0.019 | 20.57 | 69.98 | 96.25 | 99.62 |
| Smoking-related cancers (all) | 0.95 | 0.92 | 0.98 | 0.003 | 7.11 | 40.78 | 88.33 | 98.71 |
| Non-smoking related cancers |  |  |  |  |  |  |  |  |
| Prostate cancer | 0.98 | 0.95 | 1.01 | 0.234 | 77.05 | 96.80 | 99.70 | 99.97 |
| Leukemia | 0.98 | 0.96 | 0.995 | 0.012 | 34.13 | 82.35 | 98.08 | 99.81 |
| Breast cancer | 0.94 | 0.89 | 0.99 | 0.028 | 30.32 | 79.66 | 97.73 | 99.77 |
| Melanoma | 0.98 | 0.93 | 1.03 | 0.447 | 76.52 | 96.70 | 99.69 | 99.97 |
| Lymphoma | 1.01 | 0.95 | 1.07 | 0.764 | 79.89 | 97.28 | 99.75 | 99.98 |
| Ovarian cancer | 1.01 | 0.96 | 1.06 | 0.797 | 81.76 | 97.58 | 99.78 | 99.98 |
| Non-smoking-related cancers (all) | 0.98 | 0.97 | 0.995 | 0.009 | 34.62 | 82.65 | 98.13 | 99.81 |
| Smoking/non-smoking related cancers |  |  |  |  |  |  |  |  |
| All cancers | 0.98 | 0.96 | 0.99 | 0.000 | 2.19 | 16.74 | 68.86 | 95.71 |

**Figure 3.**
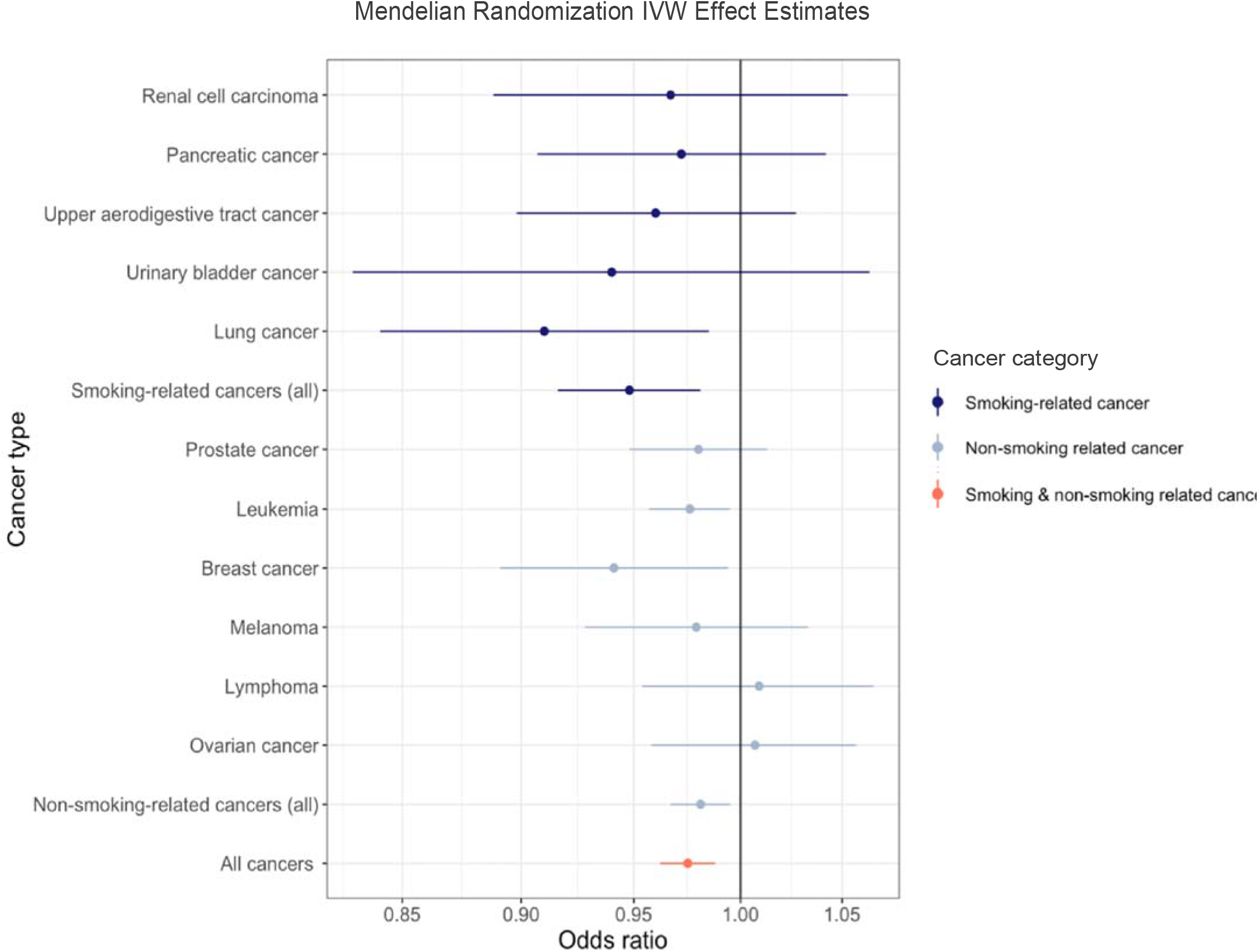
Odds ratio of Alzheimer’s disease per genetically predicted increase in risk of cancer. Each circle represents the inverse-variance weighted Mendelian randomization estimate for the causal effect of the corresponding genetically predicted cancer on Alzheimer’s disease. Dark blue represents smoking-related cancers, steel blue represents non-smoking related cancers, and orange represents all cancers in aggregate. Approximately 86% of points fell to the left of the null (OR=1), while 14 % fell to the right of the null. Genetically predicted lung cancer, leukemia, and breast cancer, as well as smoking-related cancers, non-smoking related cancers, and all cancers together were associated with significantly lower odds of Alzheimer’s disease.

Genetically predicted non-smoking related cancers were associated with 1.9% lower odds of AD (OR 0.98, 95% CI 0.97 to 0.995, p=0.0091; BFDR_50%_=34.62%, BFDR_10%_= 82.65%, BFDR_1%_=98.13%, BFDR_0.1%_=99.81%) per 1-unit higher log odds of cancer (Table 2 and Figure 3). Within non-smoking related cancers, genetically predicted prostate cancer, melanoma, lymphoma, and ovarian cancer were not associated with differential odds of developing AD. However, genetically predicted leukemia was associated with 2.4% reduced odds of AD (OR 0.98, 95% CI 0.96 to 0.995, p=0.012; BFDR_50%_=34.13%, BFDR_10%_=82.34%, BFDR_1%_=98.09%, BFDR_0.1%_=99.81%) per 1-unit higher log odds of cancer, and breast cancer was associated with 5.9% lower odds of AD (OR 0.94, 95% CI 0.89 to 0.99, p=0.028; BFDR_50%_=30.32%, BFDR_10%_=79.66%, BFDR_1%_=97.73%, BFDR_0.1%_=99.77%) per 1-unit higher log odds of cancer (Table 2 and Figure 3).

When genetic predictors of cancers at all sites under consideration were pooled, cancer was associated with a 2.5% reduction in odds of AD (OR 0.98, 95% CI 0.96 to 0.99, p=0.00027; BFDR_50%_=2.18%, BFDR_10%_=16.74%, BFDR_1%_=68.86%, BFDR_0.1%_= 95.71%) per 1-unit higher log odds of cancer (Table 2 and Figure 3).

#### 3.3.2 Sensitivity analyses

##### 3.3.2.1 MR-Egger intercept test

Pleiotropy was assessed based on the intercept of MR-Egger analysis. In all cases, there was no evidence to reject the null hypothesis of no unmeasured directional pleiotropy of the genetic variants (i.e. MR-Egger intercept of 0) (Table 3).

**Table 3.** MR-Egger intercept test for unbalanced horizontal pleiotropy. In all cases, there is no evidence of unbalanced horizontal pleiotropy.

| Cancer type | Intercept | Standard error | P-value |
| --- | --- | --- | --- |
| Lung cancer | 0.024 | 0.014 | 0.118 |
| Leukemia | -0.004 | 0.008 | 0.645 |
| Breast cancer | -0.001 | 0.004 | 0.770 |
| Smoking-related cancers | -0.003 | 0.008 | 0.695 |
| Non-smoking related cancers | -0.002 | 0.002 | 0.254 |
| All cancers | -0.003 | 0.002 | 0.167 |

##### 3.3.2.2 Leave-one-out analysis

To further examine the stability of the Mendelian randomization effect estimates and identify any important outliers, each SNP was sequentially removed from the analysis, conducting Y analyses with Y-1 datapoints. The precision and direction of the association between cancer and risk of AD remained largely unaltered with this approach. Only four SNPs had a marginal influence on the overall estimate (namely, for the association between lung cancer and AD and breast cancer and AD). Plots of leave-one-out analysis showing the influence of individual SNPs on the overall effect estimate for each cancer type and risk of AD are provided in Appendix 2.

##### 3.3.2.3 Funnel plots of inverse standard error

Any heterogeneity of effect estimates, and thereby the likelihood of directional pleiotropy of SNPs, was assessed on the basis of funnel plots depicting the inverse standard error of the causal estimate for each genetic variant. Asymmetry about the vertical line is indicative of heterogeneity, possibly due to pleiotropy. Figure 4 depicts the funnel plots for lung cancer, smoking-related cancers, leukemia, breast cancer, non-smoking related cancers, and all cancers.

**Figure 4.**
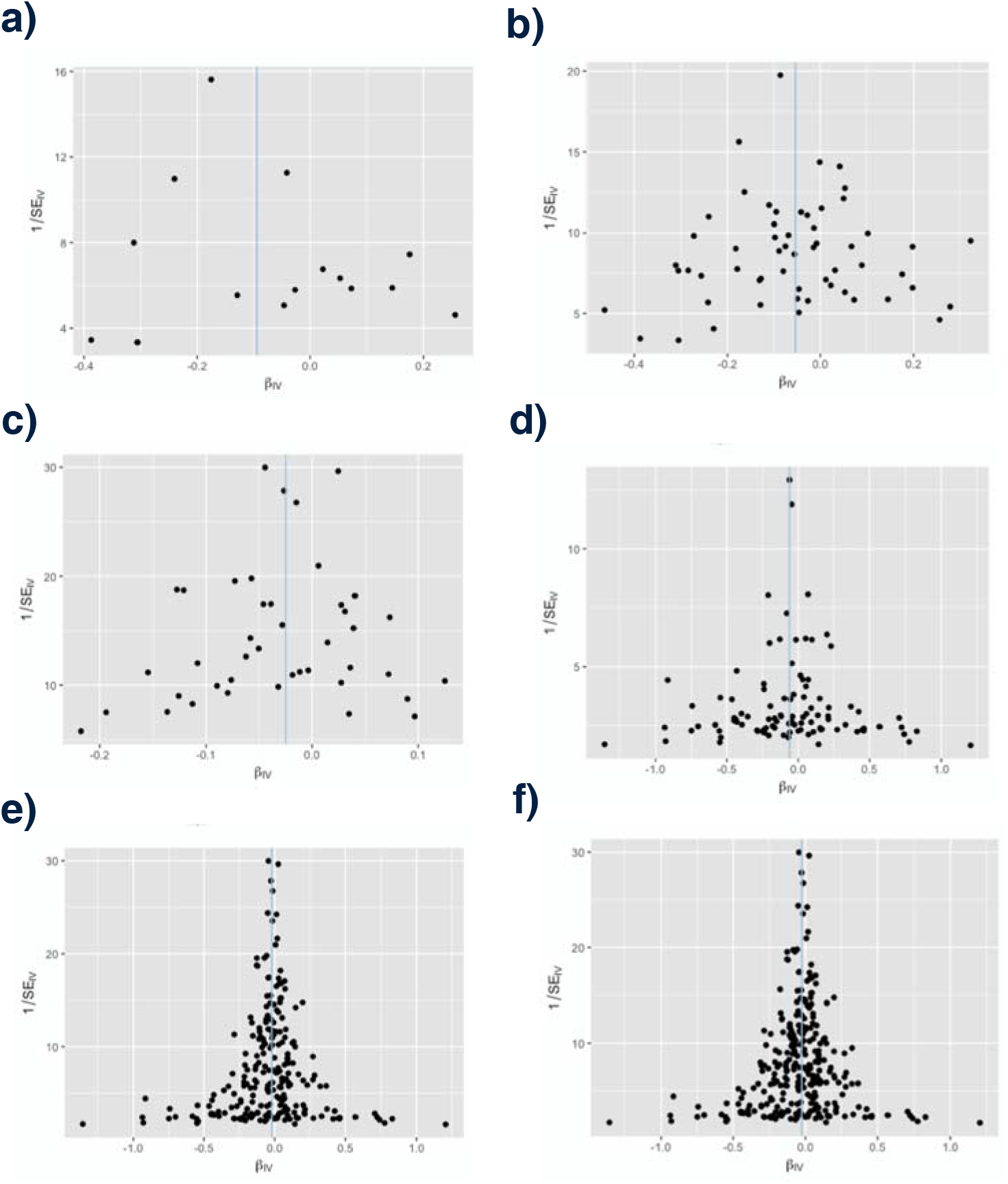
Funnel plots depicting the relationship between the causal effect of cancer on Alzheimer’s disease estimated by each genetic variant against the inverse standard error of the causal estimate. The x-axis represents the effect estimate (beta-coefficient) for risk of Alzheimer’s disease for each SNP. The y-axis represents the standard error of the estimated effect. The blue vertical line represents the inverse-variance weighted Mendelian randomization estimate using all SNPs. In all cases, the estimated effects scatter roughly symmetrically about the overall Mendelian randomization estimate, indicating precision in the estimated effects as well as absence of pleiotropic effects. a) lung cancer b) smoking-related cancers c) leukemia d) breast cancer e) non-smoking related cancers f) all cancers.estimated effects as well as absence of pleiotropic effects. a) lung cancer b) smoking-related cancers c) leukemia d) breast cancer e) non-smoking related cancers f) all cancers.

## 4. Discussion

Our results are consistent with a protective effect of some cancer-related gene variants against Alzheimer’s disease, and support genetically mediated mechanisms for the inverse epidemiological association. Specifically, genes associated with increased risk of cancer at any site were associated with a reduced risk of Alzheimer’s disease, and across individual cancer types, lung cancer, breast cancer, and leukemia were found to be associated with a statistically significant lower risk of AD. Finally, genetically predicted smoking-related cancers were more strongly associated with AD than non-smoking related cancers. This Mendelian randomization approach provides critical information on causality, beyond the former epidemiological observations, with important implications for public health and the prevention of both cancer and AD.

Taken together, the results of our Mendelian randomization complement and extend existing observational evidence of an inverse relationship between cancer and AD. Two studies synthesized observational data covering multiple cancer types to compare the relationship between smoking-related cancers and AD with the relationship between non-smoking related cancers and AD.^6 11^ In line with their findings, our study demonstrates an inverse causal relationship between genetically predicted cancers and risk of AD and, furthermore, supports a stronger relationship between genetically predicted smoking-related cancers and AD than genetically predicted non-smoking related cancers and AD.

Strengths of this study include the use of data from large GWAS of cancer, as well as the MR design, which reduces confounding and bias from reverse-causality. Furthermore, the validity of MR relies on the assumptions that the instrumental variables are strongly associated with the risk factor of interest, are not associated with any confounders, and do not influence the outcome through alternative causal pathways (Figure 5).^32^ These factors were addressed in the study design.^33^ While the lack of classical instrumental variables analysis means that instrumental variable strengths cannot be formally evaluated through the generation of an F-statistic,^34^ any instrumental bias would, in this case, be in the direction of the null, given that analyses were undertaken in separate settings for the exposure and outcome.^35^ Therefore, bias from weak instruments cannot explain the observed relationship between cancer and AD. Furthermore, while the wide range of risk factors for AD and the complex, interlaced nature of physiological signaling complicate this assumption, results of the MR-egger intercept test, as discussed above, demonstrate that pleiotropy is unlikely to explain the observed association between cancer and AD.

**Figure 5.**
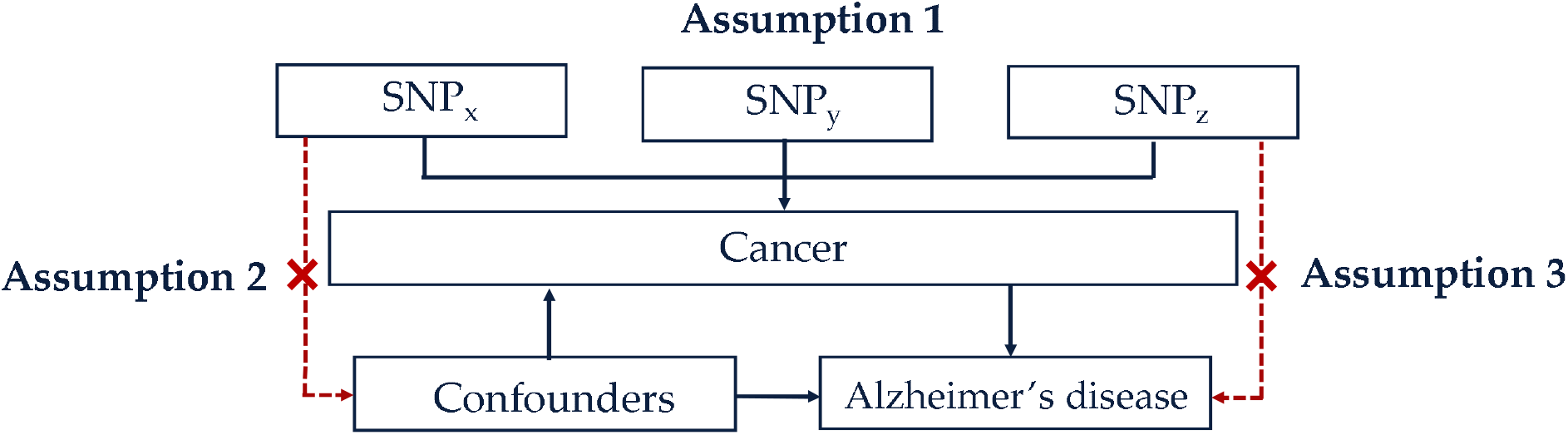
Core assumptions of Mendelian randomization. The validity of Mendelian randomization relies on three core assumptions: (1) the genetic variants are associated with the exposure (cancer), (2) the genetic variants are not associated with any confounders, and (3) the genetic variants influence the risk of the outcome (Alzheimer’s disease) through the exposure (cancer), and not any other pathways. SNP_x_, SNP_y_, SNP_z_=single nucleotide polymorphisms. Adapted from Larsson et al., 2017

Nonetheless, our results must be considered in the context of several important limitations. While summary statistics are provided in IGAP, there is a lack of covariate information, which impedes the incorporation of important factors, such as APOE allele information, education, smoking, obesity, and socioeconomic status, that can influence vulnerability to AD. Another potential limitation is the variability in the diagnostic criteria used to identify AD in IGAP studies and the inevitable possibility of misclassification bias.

We present largely concordant results from Bayesian and frequentist statistical analyses, while recognizing the very different nature of the statistical tests. The Bayesian analysis quantifies the relative probability of alternative hypotheses (including a null hypothesis) given the observed data distributions. In view of the consistency of existing observational evidence for smoking-related cancers, a BFDR prior of 50% was used, while a more stringent prior of 10% was used to evaluate non-smoking related cancers. Under these assumptions, a caveat is necessary for the interpretation of the apparent associations for breast cancer, leukemia, and non-smoking related cancers (noting the BFDR= 30% for breast cancer, 34% for leukemia, and 35% for non-smoking related cancers). However, when all cancers are considered in aggregate, or just smoking-associated cancers, the Mendelian randomization approach indicates that genes associated with cancer are associated with lower risk of AD. Twenty-eight genetic variants linked to cancer were associated with lower odds of AD, 13 of which are located either on or near genes that have been previously linked with AD (Supplementary Table 6). The remaining 15 SNPs merit further study, as they may contribute in developing novel diagnostic and prognostic tools for AD, as well as effective treatments strategies for the condition.

Expansion of this study and replication in other ethnic cohorts will be necessary to determine the external validity of our findings. Future studies should additionally seek to investigate the relationship between cancer and other forms of neurodegeneration, broadly. Ultimately, these results may provide grounds for cautious optimism about the prospects for drug repurposing from cancer to AD—which may help to shorten the timelines for dementia drug discovery—and emphasize that genetic studies can help to deconvolute the complex interrelation between these two disorders.

## Contributors

SS analyzed the data, drew the figures, and wrote the first draft of the manuscript. SS and PDP were involved in the conception and design of the study. All authors contributed to the interpretation of the results and revision of the manuscript and approved the final version. For the International Genomics of Alzheimer’s Project, the investigators contributed to the design and implementation of their study and/or provided data, but did not participate in the analysis or writing of the present study.

